# Eco-evolutionary dynamics of *Plasmodium* genotypes under mass drug administration

**DOI:** 10.1101/818039

**Authors:** Maria Bargués-Ribera, R. Guy Reeves, Chaitanya S. Gokhale

**Affiliations:** Research Group for Theoretical Models of Eco-evolutionary Dynamics, Department of Evolutionary Theory, Max Planck Institute for Evolutionary Biology, August Thienemann Str. 2, 24306, Plön, Germany; Department of Evolutionary Genetics, Max Planck Institute for Evolutionary Biology, August Thienemann Str. 2, 24306, Plön, Germany

## Abstract

Mass Drug Administration (MDA) is regarded as a potential strategy for locally interrupting transmission of human malaria under specific circumstances. However, insights on how MDA affects the eco-evolutionary dynamics of different *Plasmodium* species are not well known. We provide a computational model where the ecologically explicit life cycle of the parasite is implemented. Since the parasite inhabits two different ecological niches – human host and the mosquito – it undergoes different selection pressures during its reproduction. We use the model to perform an evolutionary analysis of the dynamics of resistance alleles under atovaquone, chloroquine and combined atovaquone-chloroquine drug treatments. Our study shows how the reduced viability of resistant parasites in the mosquito affects the spread of resistance and transmission interruption in treated human populations. Overall, results confirm that the disadvantage of drug-resistant genotypes in the mosquito vector is a good tool to achieve malaria control goals under MDA programmes.

**Author summary:** Every year there are millions of new malaria cases reported worldwide. The cause of the disease is the infection by *Plasmodium*, a protozoan which is transmitted between humans through the bite of a mosquito. Antimalarials have existed since long, but *Plasmodium* has evolved resistance to the treatment, making it necessary to develop new strategies to heal the infected humans. Lately, it has been pointed out that mosquitoes could be our allies when using drugs such as atovaquone, which resistant parasites have difficulties to reproduce in the mosquito. Here we study the scenarios in which these drugs, used in Mass Drug Administration (MDA) programmes, can interrupt the transmission of malaria in local treated populations.

## Introduction

Mass Drug Administration (MDA) is currently considered by the World Health Organisation as a potential strategy for locally interrupting transmission of human malaria in isolated low transmission areas [1]. Research studies show that resistant strains of the pathogen have a reproductive disadvantage in mosquitoes: mutant zygotes have low viability and often cannot complete their life cycle. Here we analyse how MDA programmes can help transmission interruption by reducing the parasite population size by means of both the drug effect in the humans and the interruption of the life cycle of resistant strains in the mosquito.

Anti-malarial programmes which include MDA entail simultaneously providing a substantial fraction of a human population (> 70 − 80%) with courses of drugs at repeated intervals to eliminate malaria transmission in an area. Global applications of MDA to control neglected tropical diseases like onchocerciasis, schistosomiasis and lymphatic filariasis have led to major successes on regional scales [2]. However, defining the impact of MDA in malaria control is hampered by limited data. The circumstances in which MDA is most likely to prove effective in achieving interrupted transmission are exacting and include factors such as adequate access to medical facilities, effective mosquito control, limited potential for reintroduction and relatively low levels of transmission [3]. As detailed in a recent article [4] MDA, in combination with vector control and improved surveillance can interrupt transmission for up to 6 months. MDA with vector control has also interrupted malaria transmission for sustained periods among isolated island populations. [5].

From the evolutionary perspective, intuitive Darwinian principles could reason that if alleles conferring resistance to drugs are already segregating in *Plasmodium* populations, their frequency will tend to increase during MDA programs that employ those drugs. The rise of resistance acts to reduce the efficacy of drugs, both in terms of treatments and prophylactic impact. However, there is no evidence that MDA strategies increase the probability of resistance alleles arising [6], beyond that resulting from other strategies of drug use.

Recent observational and experimental studies indicate that some drug resistance alleles encountered in *Plasmodia* interfere with the completion of its life-cycle, especially the alleles for atovaquone resistance [7]. This observation has led to the proposal that such a disadvantage could be exploited to reduce the frequency of resistant genotypes ensuring that drugs in use remain effective. The consequences that this phenomenon could have on transmission rates have not been quantified.

Herein, we propose a model that can disentangle the effects of selection in plasmodia in both the mosquito and as a consequence of human drug administration. Different to previous models which tackle resistance spread focusing exclusively in the human host [10–12], our model analyses the role of the vector in parasite dynamics. In doing so, we focus on the drug combination of chloroquine and atovaquone-proguanil (termed simply atovaquone). This particular drug combination is chosen because, (1) estimates of mosquito viability for drug resistant alleles are available for both drugs [7, 8], (2) both drugs are inexpensive and free of intellectual property claims, and (3) this combination of drugs has been reportedly used in patients without incident [13]. In general, the model applies to any drugs with similar properties.

Atovaquone’s mode of action is attributed to drug activity against the liver and pre-liver stage parasites and targets the cytochrome b protein (cytB) [14, 15]. The mitochondrial genome encodes the cytB gene and a single point mutation confers a high level of drug resistance [16]. However, the parasites which carry this resistant allele are reported to be unviable in the sexual stage of the parasite occurring in the mosquito, resulting in a promising mechanism for resistance management [7, 17]. Chloroquine acts during the erythrocytic growth of the parasite [18, 19]. Mutations of the nuclear-encoded Chloroquine Resistance Transporter gene (PfCRT) play a significant role in the widespread global resistance to this drug and other members of the antifolate class [20]. Parasites with the resistant allele K76T have been reported to have an up to 9-fold fitness reduction compared to the wild-type in the mosquito life cycle stage [8, 21]. This disadvantage, however, has not stopped the spread of chloroquine resistance in the field [22–24].

Using our model, we compare the effects of three drug regimes: atovaquone, chloroquine and their combination. We study populations under MDA and in lower drug population coverage scenarios. *Plasmodium* genotype frequencies, population size and life cycle stage of extinction are tracked in the course of transmission events between uninfected human populations. We are interested in exploring how the single and combined drug regimes perform in (1) managing already segregating resistance genotypes and (2) reducing the number of generations required to achieve local transmission interruption reliably. Our results show that MDA is effective at interrupting the parasite transmission and especially under atovaquone or combined atovaquone-chloroquine treatments due to the viability disadvantage of the resistant parasites.

### Model

Computationally, we compartmentalise the life cycle of the parasite and implement it into a mechanistic model according to its biological stages: exoerythrocytic growth, erythrocytic growth, gamete formation and transmission, fecundation and zygote selection and sporozoite formation (Fig. 1). This allows us to separate processes in the host and the vector and apply different selective pressures for resistant and susceptible parasite in different compartments. The parasite reproduces in each compartment, in discrete time, with exponential growth and multinomial sampling. Eco-evolutionary dynamics occur at two levels: within-cycle (Fig. 2), following each stage in days as a time unit, and between multiple simultaneous cycles, referred as population transmission events (Fig. 3). After each transmission event, parasite population is updated. Details of the computational and mathematical implementation are explained in the Methods section.

**Fig 1.**
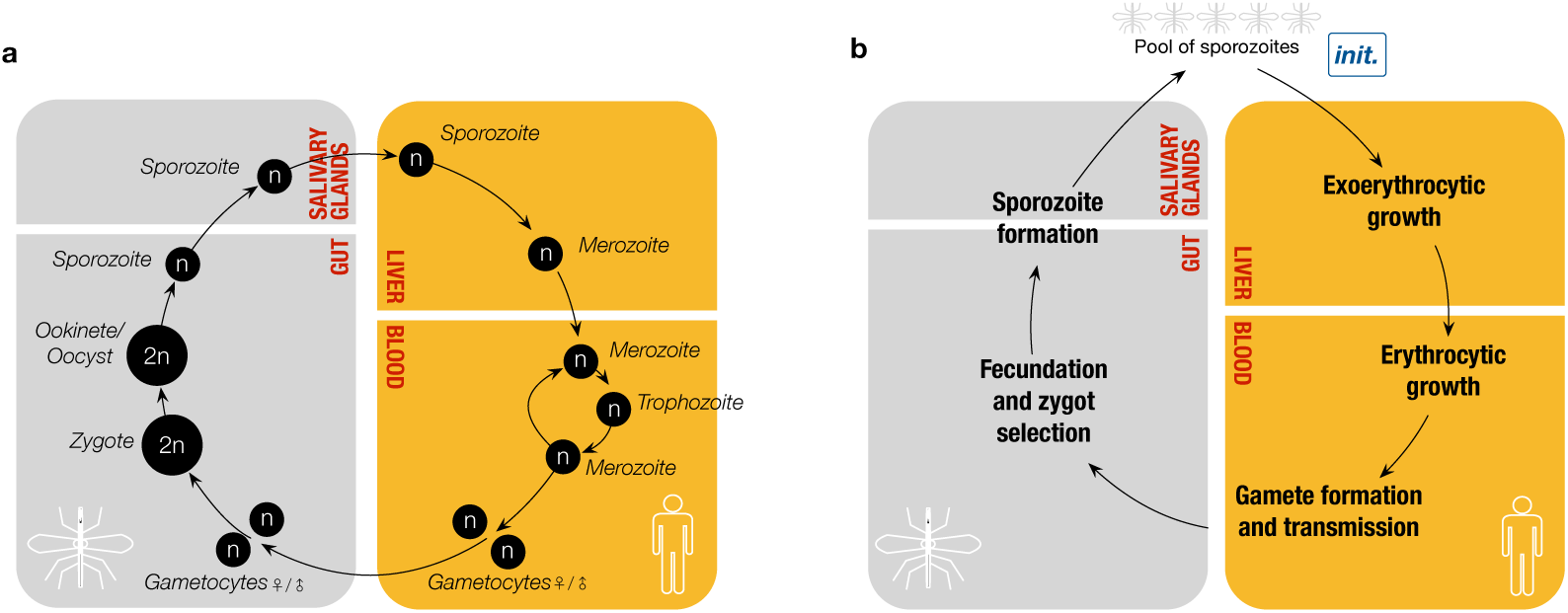
Life cycle of *Plasmodium* in human and mosquito hosts. a) Haploid (*n*) and diploid (2*n*) phases of the life cycle in human (yellow) and mosquito (grey) hosts. The mosquito bites the human and injects sporozoites, which are directed to the liver. In the liver schizonts are formed which then release merozoites. These merozoites go into the blood stream and infect erythrocytes. In the erythrocytes, merozoites mature into trophozoites and form schizonts which release merozoites again. Some of these merozoites create gametocytes, females and males in a ratio 4:1. The mosquito, when biting the infected human, takes up some gametocytes, and in its gut fecundation occurs resulting in zygote formation. The viable zygotes develop into motile ookinetes, which attach to the gut wall, mature into oocyst, and release sporozoites. These sporozoites migrate to the salivary glands, ready to be injected again in the human host in the next cycle. b) Simplified steps of life cycle in human (yellow) and mosquito (grey) hosts. For modelling purposes, we consider the life cycle starting from a pool of sporozoites which comes from the multiple mosquitoes biting humans. Then, the steps followed are: (i) exoerythrocytic growth, (ii) erythrocytic growth, (iii) gamete formation and transmission, (iv) fecundation and zygote selection, and (v) sporozoite formation. After these phases, the sporozoites formed become part of the pool of sporozoites and the cycle starts again.

**Fig 2.**
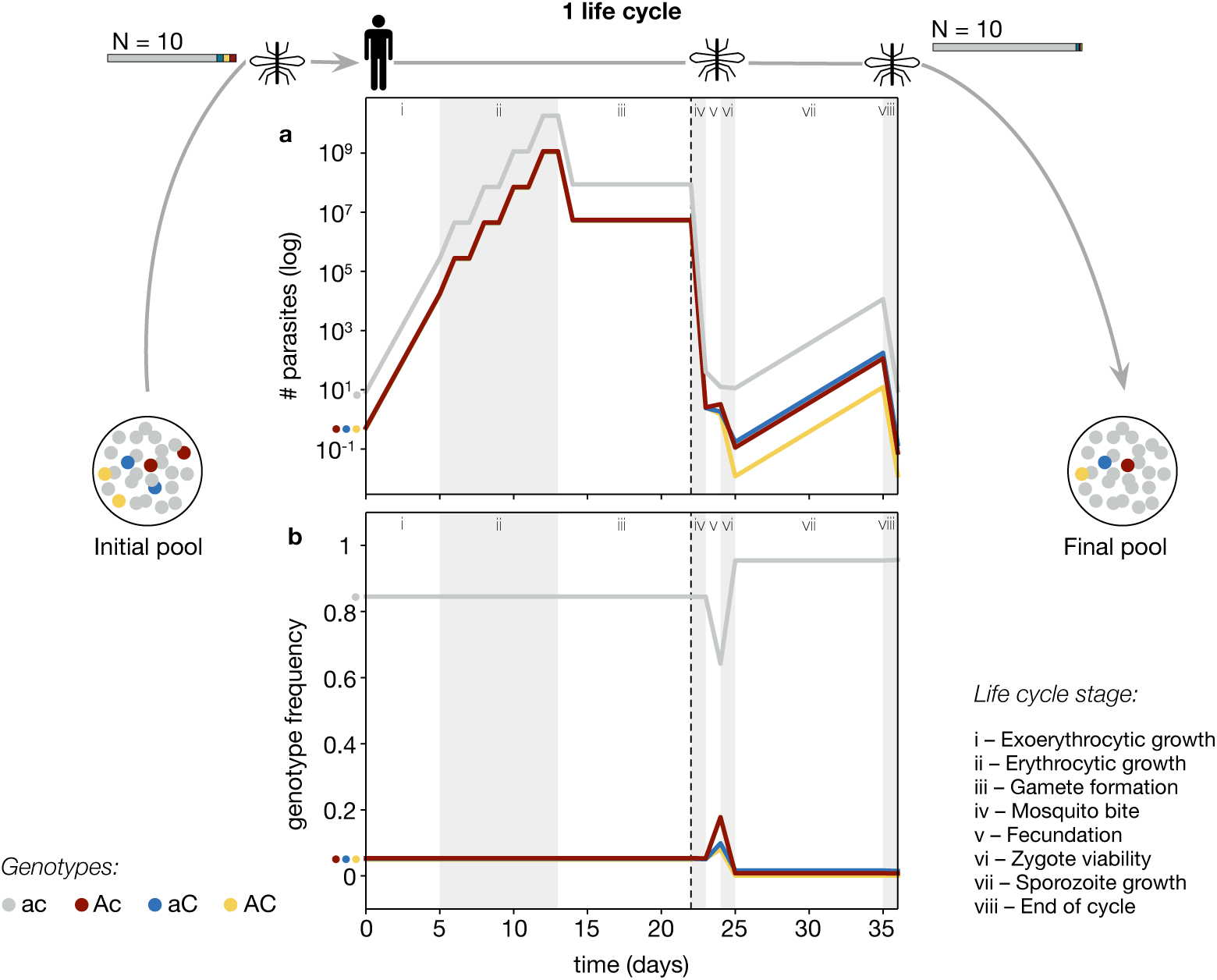
Eco-evolutionary dynamics within one life cycle without drug selection (mean of 100 realisations). From the initial pool, a mosquito carrying a sample of *N* = 10 parasites bites the human, starting the life cycle. Parasites follow the *i* − *iii* life stages in the human host and *iv* − *viii* in the mosquito (dotted line separates human-mosquito transition). At the end, after 36 days, the pool is updated with the outcome of the cycle, which corresponds to *N* = 10 unless there is extinction. a) Number of parasites according to genotype – ac (grey), Ac (red), aC (blue), AC (yellow) – during the life cycle, in log scale. b) Genotype frequencies during the life cycle.

**Fig 3.**
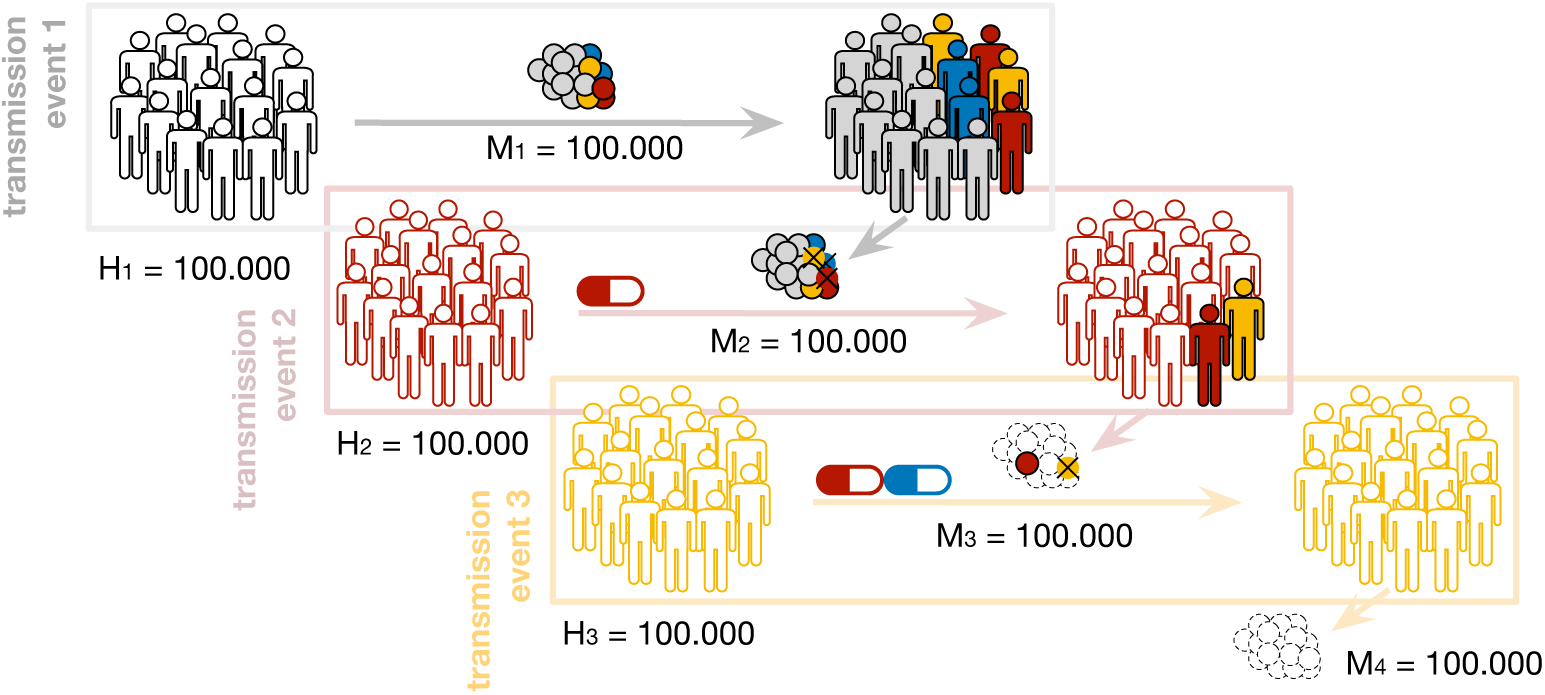
Infection of new human populations in the course of transmission events. In the first transmission event, the initial pool of plasmodia is carried by a population of mosquitoes *M*_1_. Each mosquito bites a human from a naive human population *H*_1_, which gets infected. The parasites finish their life cycle when a second population of mosquitoes *M*_2_ bites the infected humans, with some parasite extinctions due to low viability. In a second transmission event, *M*_2_ bites a new human population *H*_2_. In this example, *H*_2_ has been treated with atovaquone, which prevents some of the individuals from sustaining infection. A third population of mosquitoes *M*_3_ bites *H*_3_: the mosquitoes which bite non-infected humans become non-carriers, and extinctions due to low viability may occur. In this example, none of the individuals are bitten with a double-resistant parasite, thus none gets infected. The last population of mosquitoes *M*_4_ bites an uninfected population of humans, and disease transmission is interrupted.

We assume an isolated island with a constant population of 100,000 mosquitos (no seasonality), reproducing in synchrony without overlapping generations. Each mosquito is infected with parasites in the sporozoite form, ready to be injected to human hosts. The initial sporozoite rate, percentage of female mosquitoes with infective sporozoites in their salivary glands, is assumed for simulations to be 100%. This exceeds all field estimates from even very high transmission areas which rarely peak above 15%. However, we chose this maximal value as it provides a better appreciation of the model dynamics across the full range of possible values.

We do not model resistance allele emergence but instead assume that resistance alleles for both drugs in question already exist at appreciable frequencies in the *Plasmodium* population (0.1 each). This could realistically arise where one or both of the drugs were used in the area before MDA intervention. For one of the drugs, atovaquone, resistance is conferred by an allele of the maternally inherited mitochondrial genome [7], while the second drug, chloroquine, is via mutations in the nuclear genome [20]. This dictates that during segregation of both alleles during sexual reproduction double resistant genotypes are generated at a much higher frequency than if recombination between nuclear alleles was necessary.

While the viability of resistant alleles in mosquitos is the phenomena we are most focused on elucidating the impact of, we have relaxed the intensity of the effect relative to the published estimates. This allows for the possibility that published estimates may represent the more extreme end of possible values for viability in the field. Consequently, for illustration of the model, the viability in mosquitoes of atovaquone resistant alleles is increased to 0.05 (from non-viable in [7]), while for chloroquine-resistant alleles viability is set to 0.3 (see SI).

We do make the simplifying assumption that each human bitten by mosquitoes is initially uninfected regardless of their history or whether they are subject to drug treatment. The percentage coverage of humans administered is studied with the values 25% (low), 50% (medium), 75% (high) and 100% (full), where high and full coverage are considered MDA scenarios.

Given the set of parameters and the justifications for their ranges, we proceed with the computational model. We estimate quantities of interest such as the extinction of the parasite, the rise of resistance and the impact of the drug courses when administered with different population coverage.

## Results

Ten consecutive transmission events (non-overlapping mosquito generations) are simulated for different treatment scenarios (drug regime and population coverage). After each transmission event we record the genotype frequencies (evolutionary dynamics), size of the population of parasites (ecological dynamics) and the number of extinctions in humans and mosquito vectors. We perform 100 realisations of the complete process to account for the stochasticity in the parasite lifecycle.

### Evolutionary dynamics: low levels of viability in mosquitoes in partial drug coverage prevents the spread of resistance

Results show presence of the wild-type parasite strain after ten transmission events in all treatments for low (25%), medium (50%) and high (75%) coverage; however, full (100%) drug coverage provokes the resistant strains to outcompete the sensitive strains for all drug regimes (Fig. 4). When comparing atovaquone and chloroquine, we observe that the difference in the viability value (for atovaquone, 0.05; for chloroquine, 0.3) changes the frequency of resistant strains especially in high drug coverage. For low drug coverage, resistance does not spread in any of the single drug regimes. For medium drug coverage of chloroquine, resistant strains of chloroquine remain in low frequency until the 9th transmission event; in the 10th event only sensitive strains are present. In contrast, under medium coverage of atovaquone the resistant atovaquone strains remain only until the 3rd transmission event: the antagonistic selective pressure is strong enough to prevent their further spread. High drug coverage for chloroquine provokes maintenance of resistant strains at frequencies higher than 0.4 for all transmission events after the 3rd; whereas high atovaquone coverage prevents resistance spread from the 4th event. In the scenarios of full drug coverage, resistant strains take over the parasite population from the 1st event in all drug regimes.

**Fig 4.**
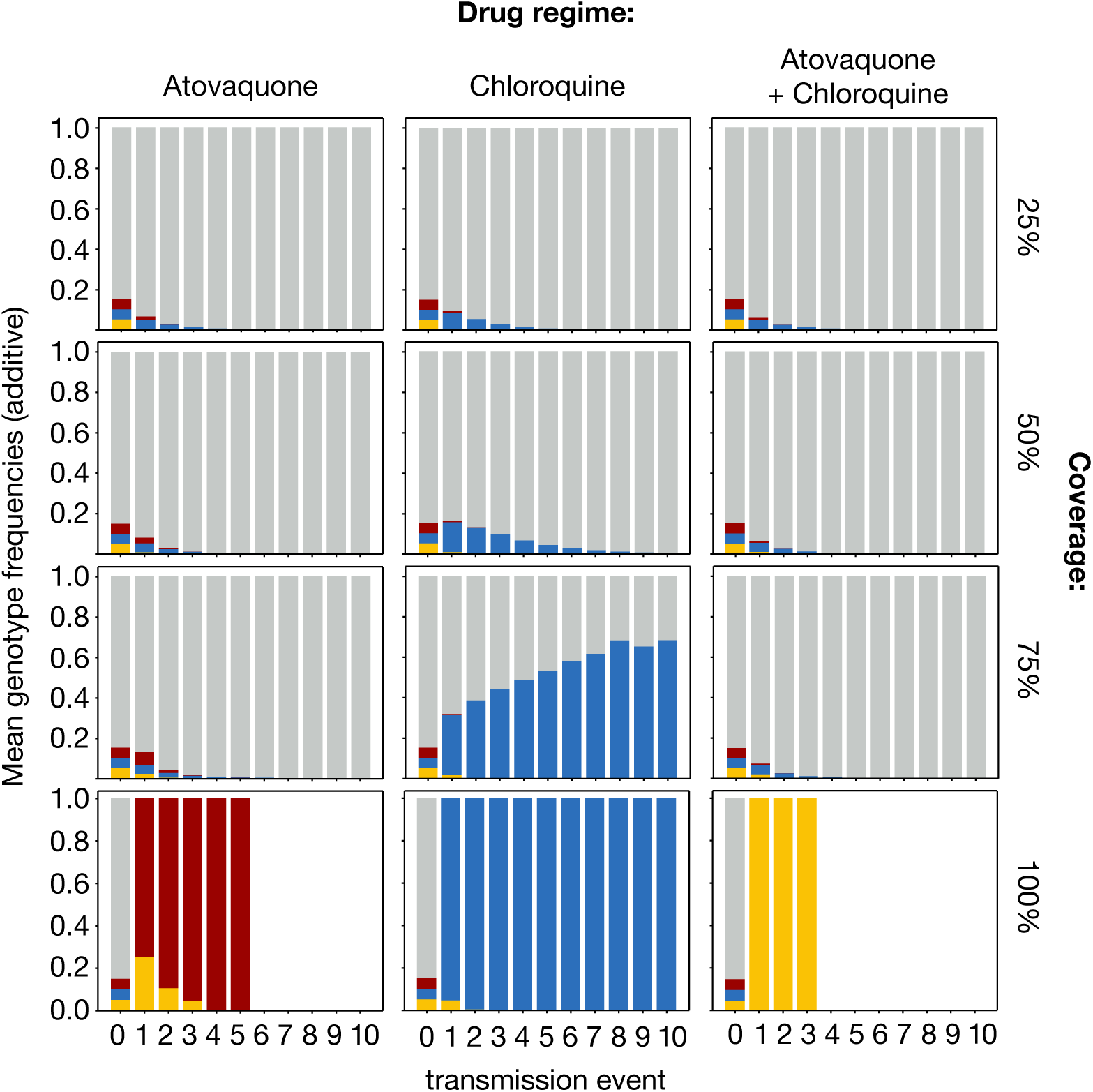
Evolutionary dynamics of the parasite pool in different drug regimes and drug population coverages (mean of 100 realisations). Mean genotype frequencies for the parasite population at the end of each transmission event are shown as cumulative bars. All drug treatment scenarios start with the same initial condition, prevalent wild-type (grey) and equal proportions of single atovaquone resistant (red), single chloroquine resistant (blue), double resistant (yellow). Dynamics are shown for three drug regimes: atovaquone, chloroquine and both atovaquone and chloroquine; in low (25%), medium (50%), high (75%) and full (100%) population coverage. The sum of all genotype mean frequencies has been normalised to 1 for all transmission events in which parasites were present. Blank events correspond to the absence of parasites.

The atovaquone-chloroquine drug combination shows qualitatively similar results to the atovaquone treatment, with an increased efficiency in interrupting parasite transmission in full drug coverage. For low, medium and high drug coverage, the resistant strains remain in very low frequencies, while the double-resistant takes over the parasite population after the 1st event when there is full coverage.

Overall, low levels of viability result in a stronger selective pressure than drug treatment under partial drug coverage, preventing the resistant strains to spread. Chloroquine resistance, which has more viability, can remain for longer in the parasite population than atovaquone resistance. In contrast, full drug coverage reverses the selection and resistant strains can outcompete their sensitive counterparts despite the mosquito disadvantage.

### Ecological dynamics: full drug coverage facilitates interruption of transmission of the parasite in single atovaquone and combined drug regimes

To interpret the results from evolutionary dynamics, we need to understand the variations in the ecological dynamics. When studying the parasite population size, we observe that the exponential decay of the pool size increases in accordance to the drug coverage and that its slope value depends on the drug regime (Fig. 5). The rate of population decay for low and medium treatment is similar for all the single and combined drug regimes: by the end of the 10th event, population size is reduced to < 10^5^ with low drug coverage and < 10^3^ for medium drug coverage.

**Fig 5.**
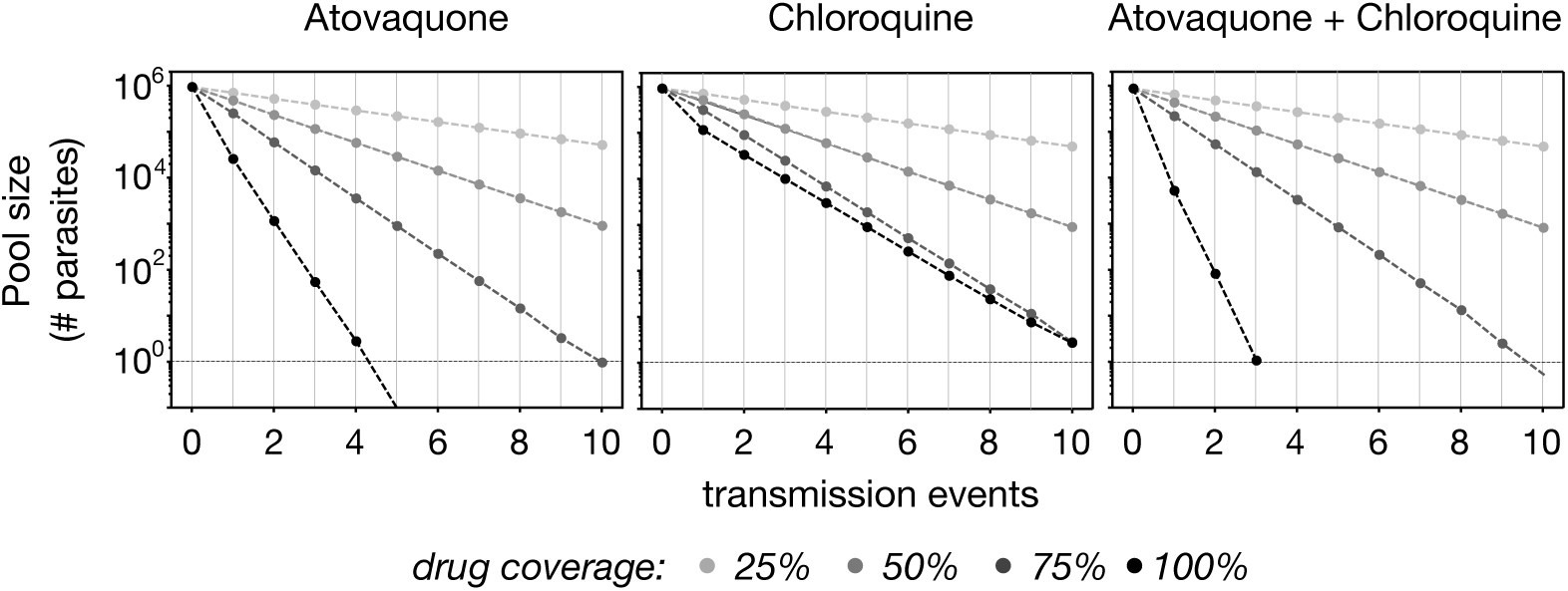
Ecological dynamics of the parasite pool in different drug regimes and drug population coverages (mean of 100 realisations). Size of the pool (as mean number of parasites, in logarithmic scale) is updated after each of the ten transmission events, depending on the drug regime – atovaquone, chloroquine and both atovaquone and chloroquine – in low (25%), medium (50%), high (75%) and full (100%) population coverage (with grey scale correspondance). Population extinction is indicated in the event in which occurs as a white dot below the threshold of population with one parasite (line).

For high population coverage, the first transmission interruption occurs for atovaquone and the combined atovaquone-chloroquine regimes (before the 9th transmission event) (Fig. 6). For chloroquine, transmission interruption happening in the 10th transmission event is the most common scenario. However, full coverage reduces the population size rapidly (Fig. 5), interrupting transmission in the 4th event for single atovaquone and the 3rd event for combined atovaquone-chloroquine treatment (Fig. 6). For chloroquine, transmission interruptions occur similarly in high and full drug coverage, being the most frequent event of interruption in full coverage (Fig. 6).

**Fig 6.**
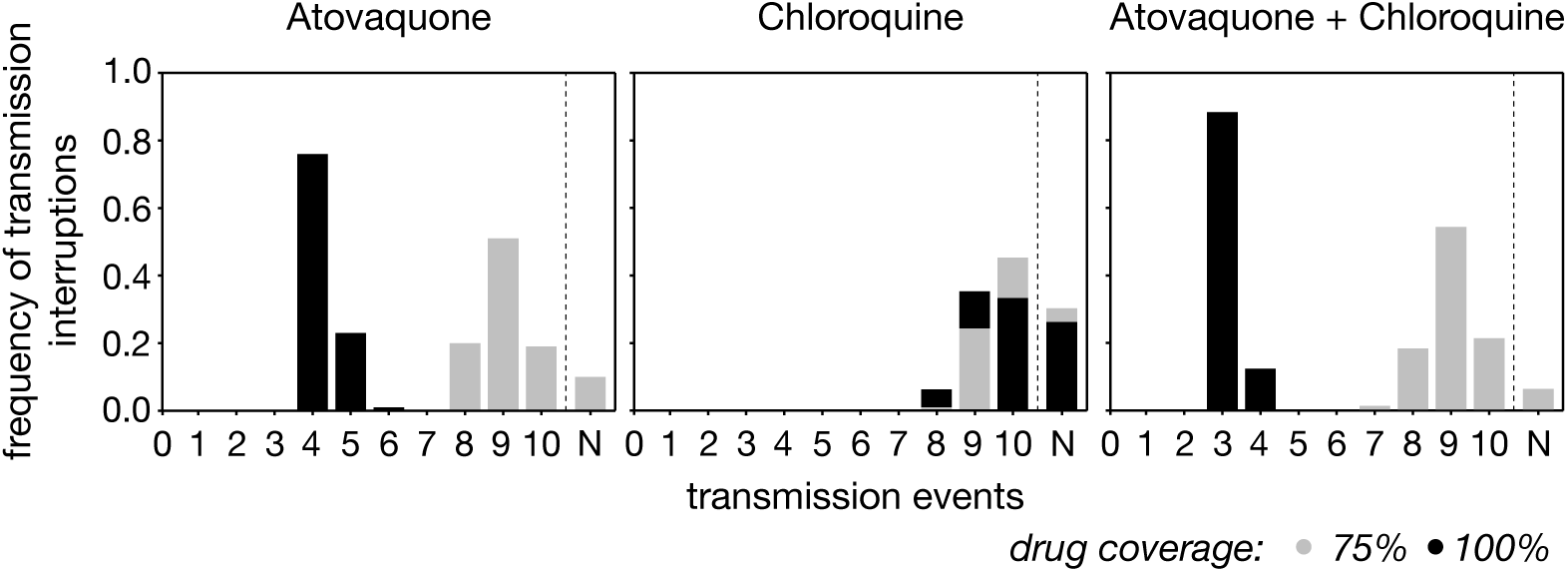
Variance in the event of transmission interruption in different drug regimes and mass drug administration scenarios (variance in 100 realisations). In one life cycle extinction of the parasite can occur in the humans or in the mosquito. Transmission interruption occurs when during one event all the parasites are eradicated in all lifecycles. A 100 realisations of this process provide a distribution of the time when such transmission interruption occurs. Distribution of the number of realisations in which transmission interruption occurs in high (grey) and full (black) drug coverages. The event number corresponds to the event in which transmission interruption occurs (i.e. no parasites at the end of the event). Transmission interruption can occur later than the 10th transmission event (event N).

In general, ecological dynamics show that there is little difference in the time of transmission interruption between the single atovaquone and the combined atovaquone-chloroquine treatment, especially when considering the variation caused by stochastic effects. Results are similar for the different drug regimes when the treatment is administrated to 75% of the population.

### Eco-evolutionary dynamics and drug efficiency: low drug coverage maintains the wild-type genotype causing drug treatment to be effective for longer

The interaction between ecological and evolutionary dynamics needs to be analysed to understand the cause of parasite extinctions. Low drug coverage relates to high numbers of parasites, mostly drug-sensitive: this leads to high numbers of parasite population extinctions in the human host. Contrarily, full drug coverage makes the population of parasites decay rapidly, with only survival of resistant strains: although in the first event several extinctions in the human occur, further extinctions happen only in the mosquito because of the low viability of selected resistant parasites. (Fig. 7)

**Fig 7.**
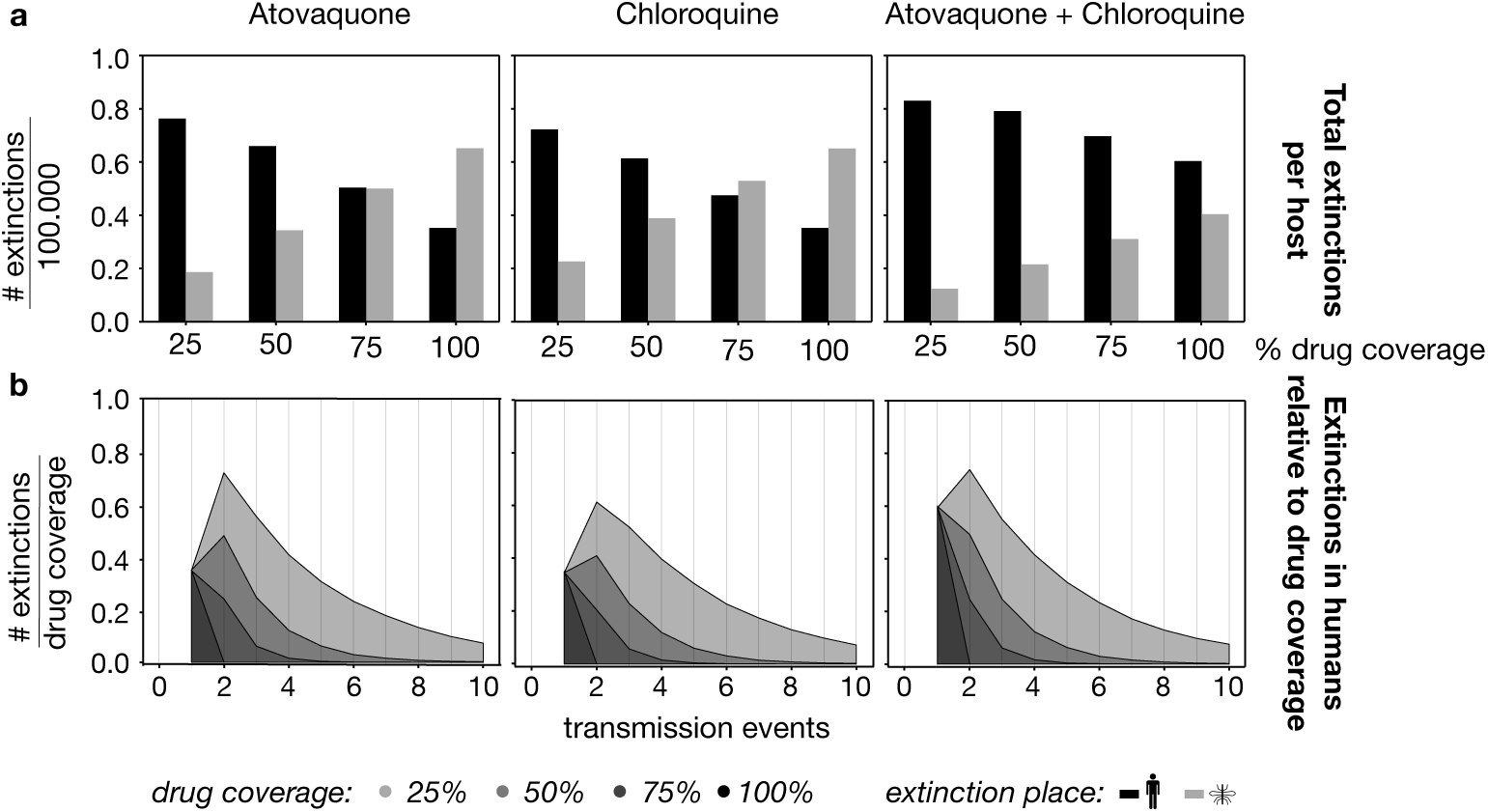
Extinctions in the parasite population in different drug regimes and drug population coverages (mean of 100 realisations). a) Mean number of extinctions that occur in the host (black) and the vector (grey) per drug treatment during all the transmission events under different drug coverage values –low (25%), medium (50%), high (75%) and full (100%)–, in bars. b) Mean number of extinctions that occur in the human host, relative to the drug coverage (in grey scale), in each transmission event.

For all drug regimes, low and medium drug coverage provoke more extinctions in the human host than in the vector, due to the high frequency of wild-type parasites. High population coverage with a single drug shows a similar number of extinctions in the two locations, with chloroquine regime showing more extinctions in the mosquito. Full coverage shows, for single drug regimes, more extinctions in the mosquito than in the human. When observing the double drug regime, instead, the number of extinctions in humans is greater than in mosquitos for all drug coverages (Fig. 7a).

From a medical perspective, the goal of drug efficiency is to prevent infection in human individuals. A proxy for efficiency would be the number of parasite extinctions in human hosts along the ten transmission events (Fig. 7b). Because of the differences in drug coverage, we observe the extinctions relative to the population that receives prophylactic treatment. The results thus indicate the utility of drug treatment in each human population. Importantly, we do not compare the total number of extinctions in humans per transmission event, but we demonstrate the relationship between extinctions and the number of treatments, i.e. efficacy.

This results show that low population coverage is, for all drug regimes, the most efficient treatment along all the events. In the first event, all drug coverages have the same efficacy because of the initial genotype frequencies, and only differing between drug regimes. The double-drug regime has more efficacy because it can eliminate more parasites than the single-drug regimes.

However, along the transmission events, the genotype frequencies change according to the evolutionary dynamics and this affects the efficacy. Drug coverage scenarios that favour the prevalence of sensitive-strains show more efficacy. Besides, the ecological dynamics also influence the treatment efficacy: when the population size is big, more humans get infected and the treatment becomes useful for more individuals. Thus, despite the similar frequencies of sensitive strain that low and high coverage of atovaquone have, treatment loses efficacy in high coverage because the low number of existing parasites. For full coverage, the treatment is very efficient in the first event, but the spread of resistant strains stops the direct effect of human healing of the treatment. This phenomena is accentuated in the double-drug regime.

## Discussion and conclusion

The finding by Goodman et al. [7, 17] that maternally inherited resistance alleles in malaria plasmodia to the drug atovaquone show limited viability in mosquitoes, led them to speculate that this could be exploited to limit the increase in frequency of such alleles in circumstances where the drug in question is widely employed, e.g. MDA. By developing and implementing our life cycle based model we have confirmed that drug resistance management is indeed feasible (Fig. 4), even in MDA programmes where ≫ 70% of the human population may be receiving the drug.

Given that globally, best practice requires the use of drug combinations for malaria treatment and prophylaxis, we chose to incorporate into our model a second drug. In doing so, we selected chloroquine, which in common with the vast majority of malaria drugs has resistance alleles encoded on the plasmodia nuclear chromosomes. Moreover, some disadvantage in the reproduction of chloroquine-resistant parasites has been reported in previous literature [8, 21].

In our model, we studied the effect of antagonistic selection pressures by including fitness parameter estimates for atovaquone and chloroquine. However, any combination of drugs with resistance alleles with suitable patterns of inheritance could be modelled across a wide range of parameter values. It is essential to note that in our modelled scenarios, resistance alleles are already initially present. The rationale for not incorporating the emergence of resistance is appropriate as there is no evidence that MDA programs increase the probability of resistance alleles arising, beyond that resulting from other patterns of drug administration [4, 6].

The insights of our MDA model relate to the change in frequencies over time of drug-resistant genotypes and impact on the goal of timely interruption of malaria transmission in an isolated population. The evolutionary findings of our model confirm the intuitive and established [25] result, that where resistance alleles are present in the parasite population, low (25%) and medium (50%) levels of population drug coverage should not act to substantially increase resistance frequency. However, at higher coverage treatment regimes ≥75% the low viability of resistance alleles for one drug in the mosquito vector can usefully manage the rise of resistance alleles for another drug.

The comparison between single and double drug treatment shows little or no synergistic interaction in combined drug use, in terms of resistance allele management. Chloroquine nuclear-encoded resistance, given by PfCRT mutation K76T (viability = 0.3), increases both in 75% and 100% treatment coverage (Fig. 4). The impact of atovaquone with the much more substantial reduction in viability (viability = 0.05) is different, as the frequency of resistance alleles only increases under full 100% coverage.

The atovaquone treatment outcomes turn out similar to the combined atovaquone-chloroquine regime, suggesting a single drug with very low viability would be sufficient for MDA programmes. This result is observed even under our conservative (but untested) assumption that the mosquito viability disadvantage is multiplicative for the double resistant plasmodia individuals.

With respects to the ecological findings of the model, we are interested in the capacity to achieve local interruption of malaria transmission. Even starting from an unrealistically high sporozoite rate of 100% where all the female mosquitoes initially are infective (1 to 2 orders of magnitude higher than observed in field studies in high transmission areas), eliminating transmission is still possible where a large proportion of the human population consents to continuous drug treatment (Figs. 5 and 6). The transmission interruption occurs within a much smaller number of transmission events in the scenarios where a drug with a substantial reduction in resistant genotype mosquito viability is employed – i.e. more than two times faster for atovaquone than for chloroquine treatment. (Fig. 6). As with the evolutionary dynamics, there is little indication of synergy, with the combination drug treatment only marginally quicker than under atovaquone alone. This is even more noticeable when we consider the variance in the number of transmission events required for transmission interruption (Fig. 6).

The combined eco-evolutionary findings of the model provide insights into the trade-off between the speed with which transmission interruption can be achieved and the extent to which resistant genotypes, including double drug-resistant ones, rise in frequency. Were it feasible to treat 100% of the population with either atovaquone or a combination of drugs, resistant alleles would probably rise to high frequency (Fig. 4). This would not prevent the MDA driven decline in the number of plasmodia circulating in the pool (Fig. 5) nor the corresponding reduction in the number of cases of human malaria (Fig. 7). On the other hand, it would mean that medical interventions to treat patients would likely not be able to rely on the effectiveness of any of the classes of drugs employed in the MDA. Consequently, in the attempt to eliminate transmission rapidly, it would be wise to make alternative drugs available to medical services that are likely to retain their effectiveness for case treatments. If a considerably slower path to transmission interruption is sought, with a population drug coverage of 75% (Figs. 5,6), high frequency of resistance can be avoided by using atovaquone or a combination of both drugs (Fig. 4). Further strategies could be investigated using the model, such as drug cycling or purposely changing drug population coverage during the treatment programme.

While our model can confirm many of the hoped-for predictions stemming from the observation of reduced viability of drug-resistant genotypes in the mosquito vector, we do not support a substantial degree of synergy stemming from a combination of resistance alleles that share this property. Consequently, it could be reasonable that an MDA based on a single drug with the most suitable properties may be sufficient to achieve malaria control goals. However, the history of malaria control indicates it is rarely wise to rely on a single mechanism, and other factors not included in our model may limit or enhance the importance of the factors highlighted here. For example, within-human fitness costs reported for resistant parasites [26] could intensify the plasmodia killing effect in humans beyond the substantial mosquito effect described here (Fig. 7A). Not explained in our model is the fact that chloroquine resistance alleles, despite inferred viability loss in mosquitoes, remain at appreciable frequencies around the world [27]. The model can thus be extended to capture more realistic scenarios, alternatively, it might be possible that genotype-specific estimates of viability in mosquitoes are to some degree context-dependent. Scenarios include the possibility that the estimated viability disadvantages of resistance alleles may be smaller than estimated [28] or may even be selected to decrease during a long-term MDA effort.

The model, as described in this paper, follows each step of the life cycle of the *Plasmodium* using available parameter estimates [29]. We envision that the model will provide a convenient basis for further elaborations. Complex models (e.g. including epidemiology, spatio-temporal heterogeneity in drug coverage) would then move towards informing the role MDA could play in leveraging genotypic viability differences towards the goal of eliminating malaria within a generation.

## Methods

We compartmentalise the life cycle of the parasite according to its biological stages and implement it into a mechanistic computational model (Fig. 1). The parasite reproduces in each compartment, in discrete time, with exponential growth and multinomial sampling. Eco-evolutionary dynamics occur at two levels: within-cycle, following each stage in days as the time unit, and between multiple simultaneous cycles, referred as population transmission events. Importantly, all resistant genotypes are present in the population from the beginning and mutation is not considered.

### Null-model: a cycle in absence of drug selection

Host-vector transmission and parasite growth happen continuously in mixed populations of humans and mosquitoes carrying *Plasmodium*. In our model, the life cycle starts with a mosquito-to-human transmission of the parasite. Then, the cycle splits into stages, which work as sequential compartments receiving a parasite input and return an output that goes on, until the cycle ends. Genetically, we define a mtDNA haploid locus for atovaquone resistance with the resistant *A* and susceptible *a* alleles. A nDNA locus for chloroquine resistance is haploid in the asexual phase and diploid in the sexual phase. The resistant *C* and susceptible *c* alleles are for chloroquine. For those stages in which the parasite is haploid, the parasite vector contains *n*_*i*_, whereas for those in which is diploid, contains *z*_*i*_ of the possible combinations shown below: 
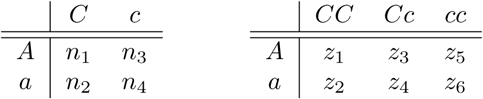

The total number of parasites as input and output for each compartment has been adjusted to the values shown in [29], in which the author reviewed and summarised quantitative data from different malaria studies. Dynamics within one cycle are shown in Fig. 2.

#### Initialisation (*t* = 0)

The cycle starts when the mosquito bites and injects sporozoites in a naive human host. A sample of *N* = 10 sporozoites is sorted by multinomial sampling according to the initial frequencies *f*_0_(*n*_*i*_) = {0.05, 0.05, 0.05, 0.85} of the *i* genotypes {*AC, aC, Ac, ac*} where the wild type is the most common genotype.

#### Exoerythrocytic growth (Day 0 to 5)

Once in the human host, sporozoites are directed into the liver, where they form schizonts and release merozoites into the blood. For sporozoites to reproduce, we use exponential growth in discrete time:

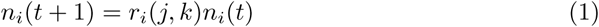

where *n*_*i*_ is number of haploid genotype *i* with *r*_*i*_(*j, k*) as its growth rate. This equation is used in all life cycle stages, that involve asexual reproduction. The growth rate of the genotype *i* depends on the life cycle stage *j* and the drug scenario *k*. Without drug selection, all genotypes *i* have the same growth within each stage *r*_*i*_(*j, k*) = *b*_*j*_. During the exoerythrocytic growth, *b*_*j*_ = 5, so each sporozoite replicates into five sporozoites per day. The population is updated until *t* = 5 [29].

#### Erythrocytic growth (Day 5 to 13)

In the erythrocytic phase, schizonts generate merozoites which infect erythrocytes in the blood. Schizonts replicate in four cycles of two days. For reproduction of the parasite in this stage, we use exponential growth (Eq. (1)). Here population is updated every two time-steps (*n*_*i*_(*t* + 2)) until *t* = 13 with a birth rate *b*_*j*_ = 16 [29].

#### Gamete formation and human-mosquito transmission (Day 13 to 23)

Gametocytes are produced with an efficiency of *ϵ* = 0.0048 [29]. Thus, the population update is the product of the current population and the efficiency rate:

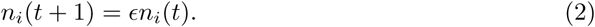

Since the population size is large, stochastic effects are ignored, and genotype frequencies do not vary. However, gametocytes are the first step of the sexual reproduction in *Plasmodium*: female and male. The female:male ratio of gametocytes is 4 : 1 [29], an important feature for fecundation. To implement sex differentiation, we multiply our population per ratio proportion:

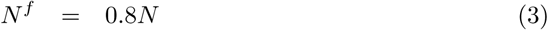

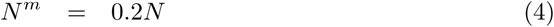

Consequently, our parasite population consists of eight types depending on sex and genotype. For these gametocytes to mate, a mosquito needs to bite the infected human and pick up both male and female gametocytes, as the fecundation takes place in the mosquito gut. The mosquito bite takes typically *N* = 48 gametocytes [29] implemented through multinomial sampling. This process introduces an ecological bottleneck for the parasite population, with drift affecting the genotype frequencies.

The parasite merozoites continue to grow and infect erythrocytes causing the disease. However, here, we keep our focus on the parasite forms that follow the complete life cycle.

#### Fecundation and zygote selection (Day 23 to 25)

In the mosquito gut, gametocytes form zygotes. Male gametocytes are the limiting factor due to the biased sex ratio. We implement sampling without replacement to match male and female gametocytes, and we obtain *Z* = *N*^*m*^ diploid zygotes, classified in six genotypes. Importantly, in the zygote genotypes, the atovaquone locus remains haploid and corresponds to the female allele, following mtDNA maternal inheritance.

The resulting zygotes *Z* are subject to viability selection. These probability of surviving are determined as *p*_*i*_ = {0.015, 0.3, 0.015, 0.3, 0.05, 1}, corresponding each value to the diploid *z*_*i*_ genotype. The values of *p*_*i*_ are set considering that (a) single atovaquone resistance (*z*_5_) has a very low probability of survival of *p*_5_ = 0.05, (b) chloroquine resistance is dominant in heterozygosis (*z*_2_ = *z*_4_, *z*_1_ = *z*_3_), (c) single chloroquine resistance (*z*_2_, *z*_4_) has a fitness disadvantage respect the wild-type (*z*_6_) of *p*_2_ = *p*_4_ = 0.3, and (d) the double resistant is assumed to have a multiplicative fitness disadvantage (*z*_1_, *z*_3_). For more details on the chosen values, see Supplementary Information.

#### Sporozoite growth and end of cycle (Day 25 to 36)

The zygotes in the mosquito gut mature into oocysts and ookinetes progressively, finally forming sporozoites and going back to haploid asexual form. In the model, the mapping from diploid to haploid follows:

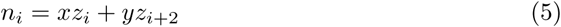

where {*x, y*} = 2 for the matching homozygote and {*x, y*} = 1 for the matching heterozygote (e.g. *n*_*AC*_ = 2*z*_*ACC*_ + *z*_*ACc*_). After the haploid conversion, the population of parasites grows exponentially following Eq. (1). For sporozoite formation, parasites reproduce with *b*_*j*_ = 2 between *t* = 25 and *t* = 35. One-fifth of the sporozoites makes it to the salivary glands (*N* = 0.2*N*), and finally, a sample of *N* = 10 parasites is randomly selected. This sample updates the parasites carried by the mosquito, thus closing the life cycle.

### Simultaneous infections in presence of drug selection and transmission events

In a population where malaria is present, there are multiple simultaneous infections. In the model, we simulate multiple simultaneous life-cycles within a transmission event. The number of simultaneous cycles is equivalent to the number of humans in the population, which in turn is equivalent to the number of mosquitoes. The entire population of parasites is conceptualised as a pool of sporozoites carried by the mosquitoes. In this pool, Plasmodia are distributed in groups of *N* = 10, representing the infected mosquitoes. Initial pool size is one million (100.000 mosquitoes carrying *N* = 10 parasites each). In absence of extinctions, pool size is maintained along the transmission event updates. During one transmission event, each mosquito bites one naive human of the population with the *N* = 10 parasites. After gametocyte formation, each human host is bitten by a new uninfected mosquito to proceed with fecundation and finish the life cycle with *N* = 10 parasites ready to be injected to a new human. In the results presented, there are ten transmission events, corresponding to approximatively one year if we consider the life cycle length of 36 days defined in this model.

#### Drug treatment

At the appropriate stages in the life cycle, we introduce atovaquone and chloroquine. Atovaquone affects the parasites before and during the liver stage, that is, during the exoerythrocytic growth. Chloroquine acts on the intra-erythrocytic parasites, that is, during erythrocytic growth. As in these stages, reproduction follows discrete exponential growth; we can include the drug as suppressed growth rate for the susceptible genotypes.

The growth rate 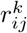 defined in Eq. 1 can be describe d now as:

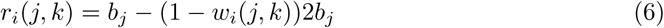

where in the presence of drugs the fitness of genotype *i* is *w*_*i*_(*j, k*) = 0.1 if the *i* is susceptible to drugs and *w*_*i*_(*j, k*) = 1 if it is resistant. Consequently, during treatment, the growth rate of susceptible genotypes becomes negative, indicating drug effectiveness. The susceptible genotypes have their fitness decreased exclusively in the stage where their nemesis drug is active, even in the case of double treatment.

#### Drug population coverage

The model allows the study of different drug population coverages by using the drug fitness disadvantage in the desired proportion of human-mosquito life cycles. The rest of life cycles are affected only by the viability selection of zygotes in the mosquito.

#### Extinctions

We consider extinction the scenario in which the initial inoculum of *N* = 10 parasites is reduced to *N* = 0, either in the human or in the mosquito. In the absence of drugs, the parasite can go extinct in the mosquito: under viability selection, all zygotes except for the wild-type genotype have a probability of survival less than 1, meaning that if by chance there is no wild-type, the whole population can die out. Also, if by stochastic means the gametocytes taken by the mosquito are from the same sex, there is no zygote formation and thus extinction. On the other hand, drug administration causes the death of the susceptible parasites in humans. If the corresponding resistant strain is present in the initial inoculum, there is no extinction in the human host.

## Acknowledgments

The authors acknowledge the generous funding from the Max Planck Society.

## Supporting information

### Assessing the levels of viability in chloroquine-resistant parasites

Chloroquine resistance is well documented, as it emerged in the late 1950’s in Asia and nowadays it affects several countries with endemic malaria. Thus, although a disadvantage has been reported for chloroquine-resistant parasites in the mosquito phase, it has not prevented the spread of resistance. Here we test different levels of viability for the parasite genotypes with the resistance mutation, to be compared in the main results with the atovaquone-resistant parasites (with a very low viability of 0.05).

The values tested in this section are the following: 0.05, 0.11, 0.3, 0.5, 1.0 (Fig. 8). The parameter range of viability is from 0 to 1; thus, in the lower range we choose 0.05 as equivalent to the atovaquone-resistance viability and we choose the upper extreme 1 in which there is no disadvantage. Because the paper [8] reports a 9-fold disadvantage of the resistant parasites, the value 0.11 is also tested. In the intermediate range, 0.3 and 0.5 are chosen arbitrarily.

**Fig 8.**
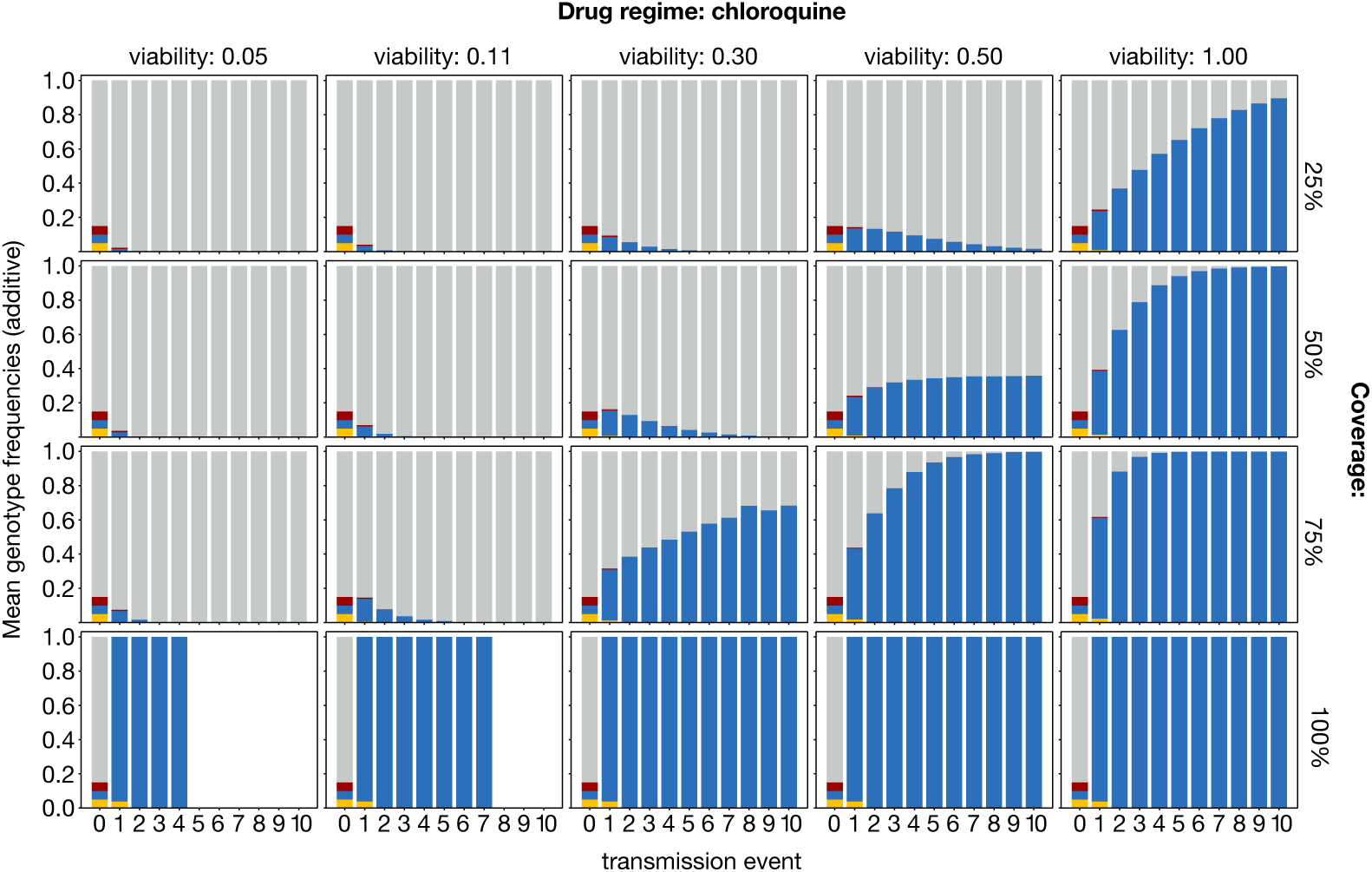
Evolutionary dynamics of parasite pool in different values of viability of chloroquine-resistant parasites (mean of 100 realisations). Mean genotype frequencies for the parasite population at the beginning of each event are shown as cumulative bars. All plots correspond to chloroquine treatment scenarios, with low (25%), medium (50%), high (75%) and full (100%) population coverage. All plots start with the initial frequencies for wild-type (grey), single atovaquone resistant (red), single chloroquine resistant (blue), double resistant (yellow). The sum of all genotype mean frequencies has been normalised to 1 for all transmission events in which parasites were present. Blank events correspond to transmission interruption.

Given the known spread of chloroquine resistance, the final choice corresponds to the smallest value which shows resistance maintenance in medium and high levels of drug coverage. The lower values 0.05 and 0.11 underestimate the viability of resistant zygotes: even in scenarios of high drug coverage (75%), resistance does not show to be prevalent from early transmission events. Disregarding the full viability of maximum value 1, which quickly allows the resistant parasites to reach high genotype frequencies, we can focus on the intermediate values: 0.3 and 0.5. Given that the prevalence of resistant chloroquine strains is usually less than 20% in countries in which drug coverage is not massive (i.e. scenarios of low or medium drug coverage), both could be realistic. However, for the focus of the study and the comparison with atovaquone-resistant parasites, we prefer not to overestimate the viability and thus choose as adequate value of study a viability of 0.3.

